# Framework for Rapid Comparison of Extracellular Vesicle Isolation Methods

**DOI:** 10.1101/2020.10.13.337881

**Authors:** Dmitry Ter-Ovanesyan, Maia Norman, Roey Lazarovits, Wendy Trieu, Ju-Hyun Lee, George M. Church, David R. Walt

## Abstract

Extracellular vesicles (EVs) are released by all cells into biofluids and hold great promise as reservoirs of disease biomarkers. One of the main challenges in studying EVs is a lack of methods to quantify EVs that are sensitive enough and can differentiate EVs from similarly sized lipoproteins and protein aggregates. We demonstrate the use of ultrasensitive, single molecule array (Simoa) assays for the quantification of EVs using three widely expressed transmembrane proteins: the tetraspanins CD9, CD63, and CD81. Using Simoa to measure these three EV markers, as well as albumin to measure protein contamination, we were able to compare the relative efficiency and purity of several commonly used EV isolation methods in plasma and cerebrospinal fluid (CSF): ultracentrifugation, precipitation, and size exclusion chromatography (SEC). We further used these assays, all on one platform, to improve SEC isolation from plasma and CSF. Our results highlight the utility of quantifying EV proteins using Simoa and provide a rapid framework for comparing and improving EV isolation methods from biofluids.

## Introduction

Extracellular vesicles (EVs) are released by all cell types and are found in biofluids such as plasma and CSF. EVs contain contents from their donor cells, providing broad non-invasive access to molecular information about cell types in the human body that are otherwise inaccessible to biopsy (1). Despite the diagnostic potential of EVs, there are several challenges that have hampered their utility as biomarkers. EVs are heterogeneous, present at low levels in clinically relevant samples, and difficult to quantify (2–4). Due to these challenges, there is a lack of consensus about the best way to isolate EVs from biofluids (5–7).

Several techniques have been used in attempts to quantify EVs. These methods, such as nanoparticle tracking analysis (NTA), dynamic light scattering (DLS), and tunable resistive pulse sensing (TRPS), aim to measure both particle size and concentration (3). A major limitation of these methods is that they cannot discriminate lipoproteins or particles of aggregated proteins from EVs (5, 8–11). In addition, they are all physical methods that provide no information about the biological nature of the particles being measured. Since biofluids, and plasma in particular, contain an abundance of lipoproteins and protein aggregates at levels higher than those of EVs (8, 12), these methods are ill-suited for quantifying EVs (2). Lipid dyes have also been used to label and measure EVs (13, 14), but these dyes also bind to lipoproteins and lack sensitivity (2). There are also numerous efforts to apply flow cytometry to the analysis of EVs, but due to the small size of EVs, obtaining quantitative measurements using this approach remains challenging (15–17).

A feature of EVs that distinguishes them from both lipoproteins and free protein aggregates is the presence of transmembrane proteins that span the phospholipid bilayer (12). The tetraspanins CD9, CD63, and CD81 are transmembrane proteins that are widely expressed and readily found on EVs, often referred to as “EV markers” (2). Although none of these proteins is present on every EV, measuring three tetraspanins should be a reliable proxy for EV abundance in many contexts. We reasoned that by using immunoassays to compare the levels of tetraspanins from a given biofluid, as well as albumin as a representative free protein, we could quantitatively compare the purity and yield of different EV isolation methods.

The most commonly used method for measuring proteins in biofluids is enzyme-linked immunosorbent assay (ELISA), but this technique lacks the sensitivity to detect low-abundance proteins (18). Single molecule array (Simoa) technology, previously developed in our lab but now commercially available, converts ELISA into a digital readout (19). Simoa assays can be orders of magnitude more sensitive than traditional ELISAs (20), which is particularly useful for EV analysis as the levels of EV proteins are often low in clinical biofluid samples (18). We have previously applied Simoa to the investigation of L1CAM, a protein thought to be a marker of neuron-derived EV, showing it is not associated with EVs in plasma and CSF (19).

In this study, we demonstrate the application of Simoa for relative EV quantification by comparing different EV isolation methods from human biofluids. In particular, we applied Simoa to compare EV isolation methods from human plasma and cerebrospinal fluid (CSF) using three of the most commonly used isolation techniques: ultracentrifugation, precipitation, and size exclusion chromatography (SEC). By also measuring levels of albumin using Simoa, we were able to determine both relative purity and yield for each technique in the same experiment. We then applied these Simoa assays to screen several parameters of SEC and develop improved EV isolation methods from plasma and CSF, demonstrating the utility of this approach for EV analysis.

## Results

### Framework for quantifying relative EV yield and purity

We set out to quantify the relative difference in yield and purity for different EV isolation methods. Starting with aliquots of the same biofluid, we reasoned that by measuring the tetraspanins CD9, CD63, and CD81 using different isolation methods, we could directly compare EV yield. By also measuring albumin, the most abundant free protein in plasma and CSF, we could compare the purity of these methods. Using single molecule array (Simoa) technology, an ultrasensitive digital ELISA, to measure all four of these proteins, we could compare EV yield and purity on one platform with high sensitivity (Figure 1a).

**Figure 1.**
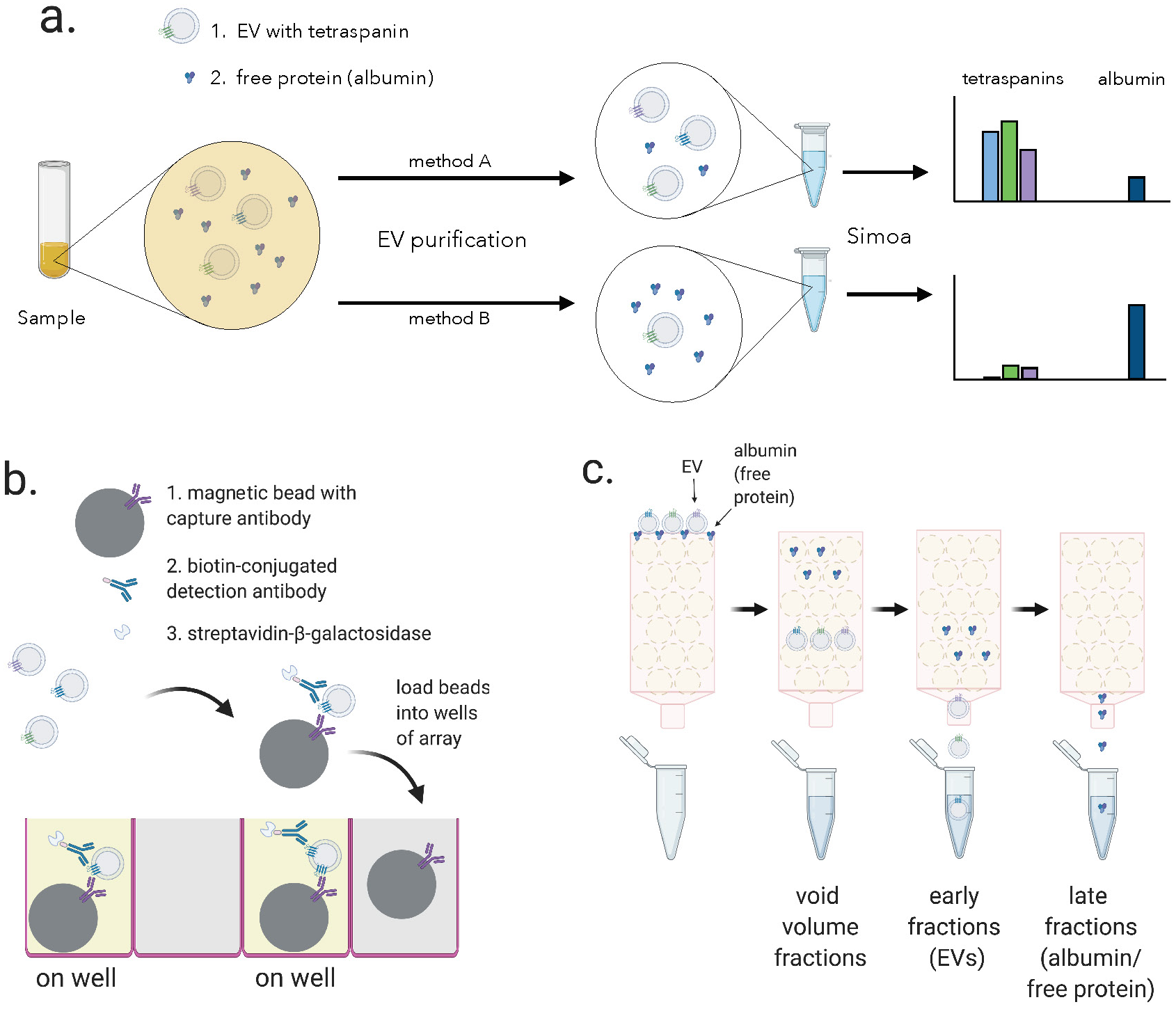
Overview of experimental framework for EV detection using Simoa and SEC. a. Different methods of EV isolation can be directly compared to assess yield and purity by measuring the three tetraspanins (CD9, CD63, CD81) and albumin. b. Single immuno-complexes are formed by binding the target tetraspanin protein on EVs to a magnetic bead conjugated to a capture antibody and a biotin-labeled detection antibody. Detection antibodies are labeled with a streptavidin-conjugated enzyme. The beads are then loaded into individual wells of a microwell array where each well matches the size of the magnetic bead limiting a maximum of one bead per well. Wells with the full immuno-complex (“on wells”) produce a fluorescent signal upon conversion of substrate, unlike wells with beads lacking the immuno-complex (“off wells”). c. EV and free proteins such as albumin in a biofluid sample are separated by size exclusion chromatography (SEC). Free proteins elute from the column in later fractions than EVs because free proteins are smaller than the pore size of the beads while EVs are larger and are excluded from entering the beads.

Although Simoa is generally used to quantify free proteins, it can also be used to analyze EV transmembrane proteins. In Simoa, unlike in traditional ELISA, individual immuno-complexes are isolated into femtoliter wells that fit only one bead per well. In a given sample, there are many more antibody-bound beads than target proteins, and therefore Poisson statistics dictate that only a single immuno-complex is present per well. This allows counting “on wells” as individual protein molecules (Figure 1b). The percentage of “on wells” can then be converted to protein concentration by comparing to a calibration curve of recombinant protein standard. We previously developed and validated Simoa assays for the proteins CD9, CD63, and CD81, showing that they are one to three orders of magnitude more sensitive than the corresponding standard ELISA assays with the same pairs of antibodies (19).

### Comparison of existing EV isolation methods

We used Simoa to directly compare three commonly used EV isolation methods from 0.5 mL samples of human plasma and CSF. For each method, we used identical samples of plasma or CSF that were pooled and aliquoted, allowing us to directly compare the different methods. To separate EVs from cells, cell debris, and large vesicles, all samples were first centrifuged and then filtered through a 0.45 μm filter. We compared ultracentrifugation (with or without a wash step), two commercial precipitation kits (ExoQuick and ExoQuick ULTRA), and two commercially available SEC columns (Izon qEVoriginal 35nm and 70nm). SEC separates EVs from free proteins based on size; proteins enter porous beads and elute from the column later than the EVs, which are much larger and less likely to enter the beads (Figure 1c). Whereas the ultracentrifugation and precipitation conditions each yielded a single sample, we collected several fractions for SEC and analyzed each fraction to assess the distribution of EVs relative to albumin.

We quantified EVs by measuring the levels of CD9, CD63, and CD81 across the different EV isolation methods in both plasma and CSF. Since we are interested in all EVs, as opposed to subsets with a specific marker, we quantified EV yield by averaging the levels of the three tetraspanins. We first used the Simoa measurement (in picomoles, determined relative to a corresponding recombinant protein standard) to calculate EV recovery for each individual marker by normalizing the level of tetraspanin in each condition to the amount of that tetraspanin in fractions 7-10 of the Izon qEV 35nm SEC column (the condition with the highest EV levels in plasma). Next, we averaged the relative tetraspanin recovery values across the three tetraspanins to calculate relative EV recovery (Figure 2).

**Figure 2.**
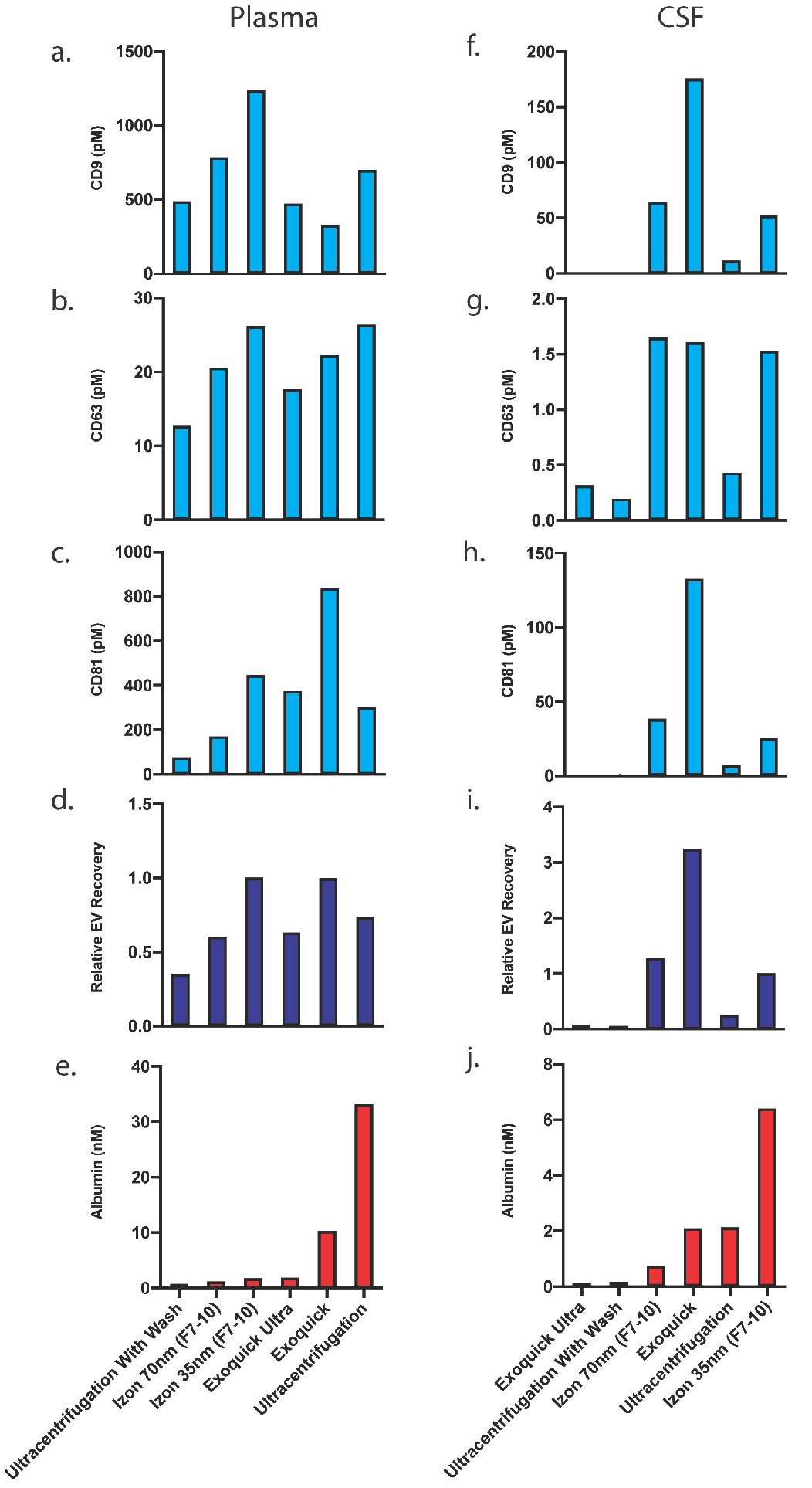
Comparison of existing methods for EV isolation in plasma and CSF. All methods are listed in order of increasing albumin levels. a-c. Individual tetraspanin yields using different isolation methods from plasma. Relative EV recoveries from plasma were calculated by first normalizing individual tetraspanin values (in pM) in each technique to those of Izon qEVoriginal 35nm EV fractions (7–10) and then averaging the three tetraspanin ratios. e. Albumin levels using different EV isolation methods from plasma. f-h. Individual tetraspanin yields using different isolation methods from CSF. i. Relative EV recoveries in CSF were calculated by first normalizing individual tetraspanin values (in pM) in each technique to those of Izon qEVoriginal 35nm fractions (7–10) and then averaging the three tetraspanin ratios. j. Albumin levels using different EV isolation methods from CSF.

After determining combined relative EV recovery and albumin concentration for each EV isolation method, we could directly compare EV recovery and purity in both plasma and CSF. In plasma, we found that the Izon qEVoriginal 35nm SEC column (collecting fractions 7-10) yielded both the highest recovery of EVs and the highest purity (ratio of EVs to albumin) of EVs (Figure 2a-e). In contrast, in CSF, ExoQuick yielded the highest recovery of EVs while Izon qEVoriginal 70nm yielded the highest purity (Figure 2f-j).

### Application of Simoa for custom SEC column optimization

Based on the superior results of commercial SEC columns, we sought to use our assays to further investigate SEC using custom columns. First, we designed an SEC stand that allows for reproducible collection of fractions and multiple columns to be run in parallel (Figure 3). We next took advantage of Simoa’s high throughput screening capability to help identify the EV-containing fractions in SEC. This enabled us to optimize EV isolation from 0.5 mL samples of plasma and CSF using SEC. We prepared our own columns to systematically test several parameters: column height (10 or 20 mL) and resin (Sepharose CL-2B, CL-4B or CL-6B).

**Figure 3:**
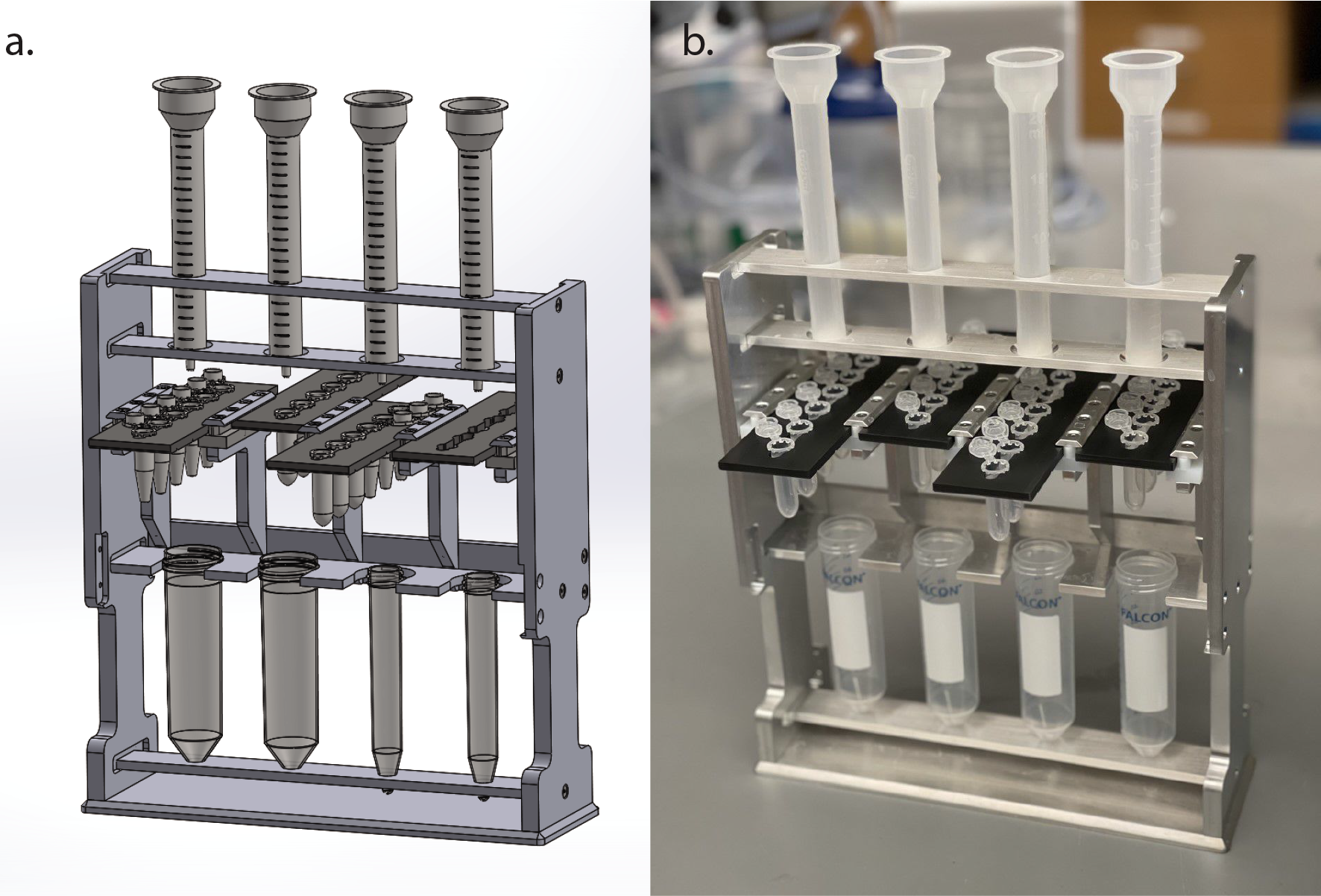
Custom stand designed for higher throughput, reproducible SEC. a. Image of SolidWorks file with custom SEC stand designed to run 4 SEC columns in parallel with “clickable” sliding collection tube holder plates that allow for easy fraction collection. b. Photograph of constructed, custom SEC stand holding four (empty) columns.

This comprehensive comparison led us to several conclusions. First, we found that resins with smaller pore sizes led to higher yields of EVs. Sepharose CL-6B, which has the smallest pore size, gave the highest yield, although it was accompanied by higher albumin contamination. For all SEC columns, higher purity could also be achieved by taking a smaller number of fractions (e.g. 7-9 instead of 7-10), albeit at the expense of lower EV yield. Additionally, we found that doubling the height of any given column from 10 to 20 mL resulted in better separation between EVs and free proteins, leading to higher purity but lower EV recovery (Figures 4 & 5). When we compared different volumes of plasma and CSF for a 10 mL Sepharose CL-6B column, we found that. as expected, larger loading volumes led to lower purity (Supplementary Figure 1).

**Figure 4:**
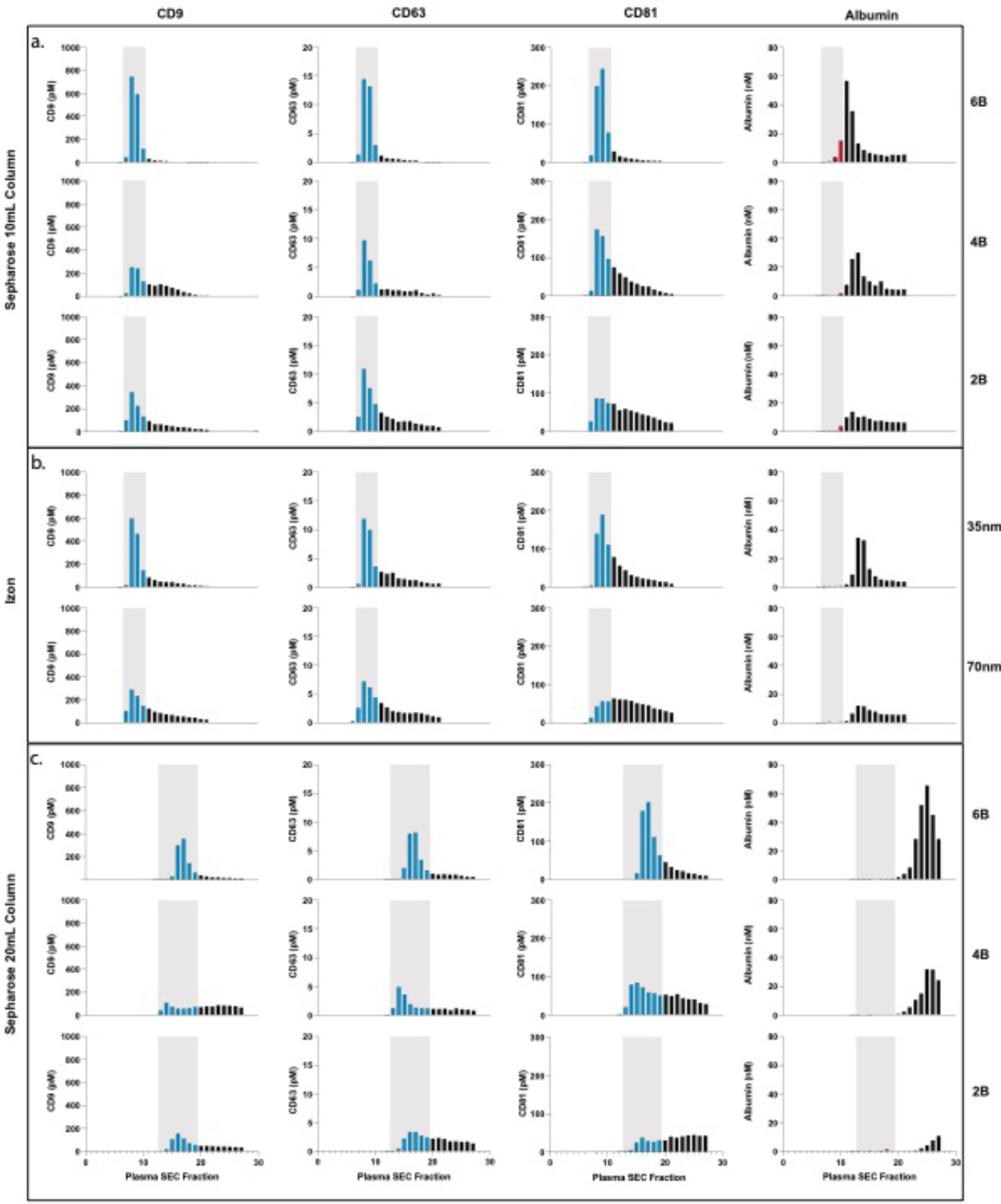
Comparison of SEC methods for EV isolation in plasma. a. Levels of tetraspanins and albumin in plasma after fractionation with 10 mL custom columns filled with: Sepharose CL-6B (top), Sepharose CL-4B (middle), and Sepharose CL-2B (bottom). b. Levels of tetraspanins and albumin in plasma after fractionation with Izon qEVoriginal 35nm column (top) and Izon qEVoriginal 70nm column (bottom). c. Levels of tetraspanins and albumin in plasma after fractionation with 20 mL custom columns; Sepharose CL-6B (top), Sepharose CL-4B (middle), Sepharose CL-2B (bottom).

**Figure 5:**
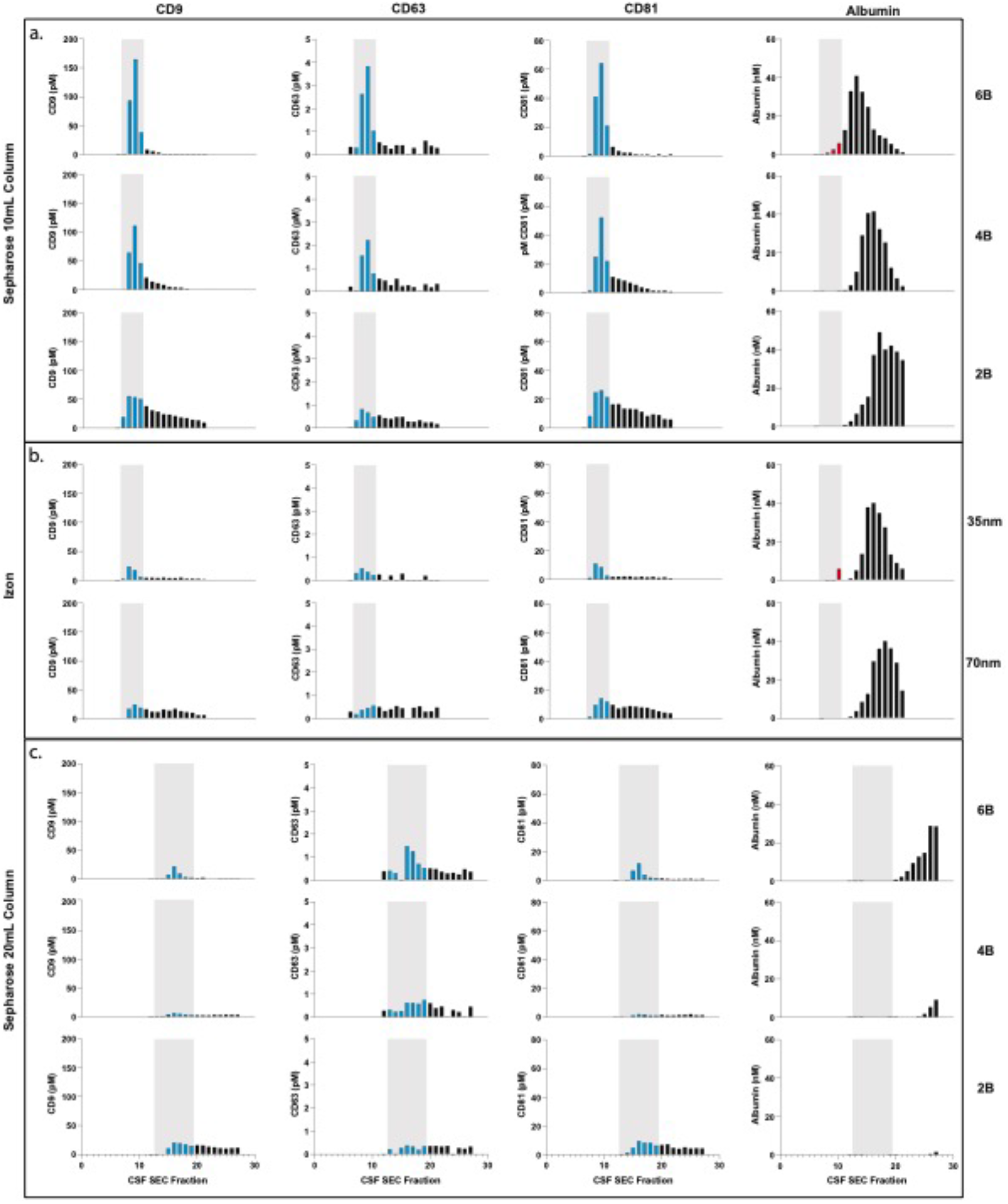
Comparison of SEC methods for EV isolation in CSF. a. Levels of tetraspanins and albumin in CSF after fractionation with 10 mL custom columns filled with: Sepharose CL-6B (top), Sepharose CL-4B (middle), and Sepharose CL-2B (bottom). b. Levels of tetraspanins and albumin in CSF after fractionation with Izon qEVoriginal 35nm column (top) and Izon qEVoriginal 70nm column (bottom). c. Levels of tetraspanins and albumin in CSF after fractionation with 20 mL custom columns; Sepharose CL-6B (top), Sepharose CL-4B (middle), Sepharose CL-2B (bottom).

### Direct comparison of custom SEC and previous methods

Combining all of the data we generated, we were able to perform a direct, quantitative comparison of the relative yields and purities of EVs across all methods tested. For both plasma (Supplementary Figure 2a) and CSF (Supplementary Figure 2b), a 10 mL Sepharose CL-6B column demonstrated the highest recovery. The 20mL Sepharose CL-4B column gave the highest purity (ratio of EVs to albumin) for plasma, while for CSF, the 10 mL Sepharose CL-4B column had higher purity than the 20 mL Sepharose CL-4B column. Although the 10 mL column had more albumin contamination in the EV fractions than the 20 mL column, the relative ratio of EVs to albumin was higher.

### Comparison of top custom SEC methods for plasma and CSF

Based on our results surveying the different SEC resins and column heights, we repeated the isolation experiments in an effort to more accurately quantify the best high yield and high purity SEC methods for plasma and CSF using another batch of biofluids with more replicates (four columns per condition). For both plasma and CSF, we compared the Sepharose CL-2B 10 mL column, used in the original SEC EV isolation publication (21) and in most subsequent SEC publications (22), to the “high yield” Sepharose CL-6B 10 mL column. We also included a Sepharose CL-4B column as the “high purity” column but, as plasma has much higher protein concentration than CSF, used 20 mL of resin for plasma and 10 mL for CSF.

Our results allow us to directly quantify the difference in EVs and albumin across these methods (Figure 6). We found that, in plasma, the Sepharose CL-6B 10 mL column provided over two-fold more EVs relative to the Sepharose CL-2B 10 mL column, but also six-fold more albumin. The Sepharose CL-4B 20 mL column, on the other hand, had similar EV levels to that of Sepharose CL-2B 10 mL column in plasma but had six-fold less albumin (Figure 6a-e), demonstrating a large increase in purity (Figure 6f). In CSF, the Sepharose CL-6B 10 mL column led to a large increase in EV yield relative to the Sepharose CL-2B 10 mL column (Figure 6g-k), but the Sepharose CL-4B 10 mL column did not lead to improved purity (Figure 6l).

**Figure 6.**
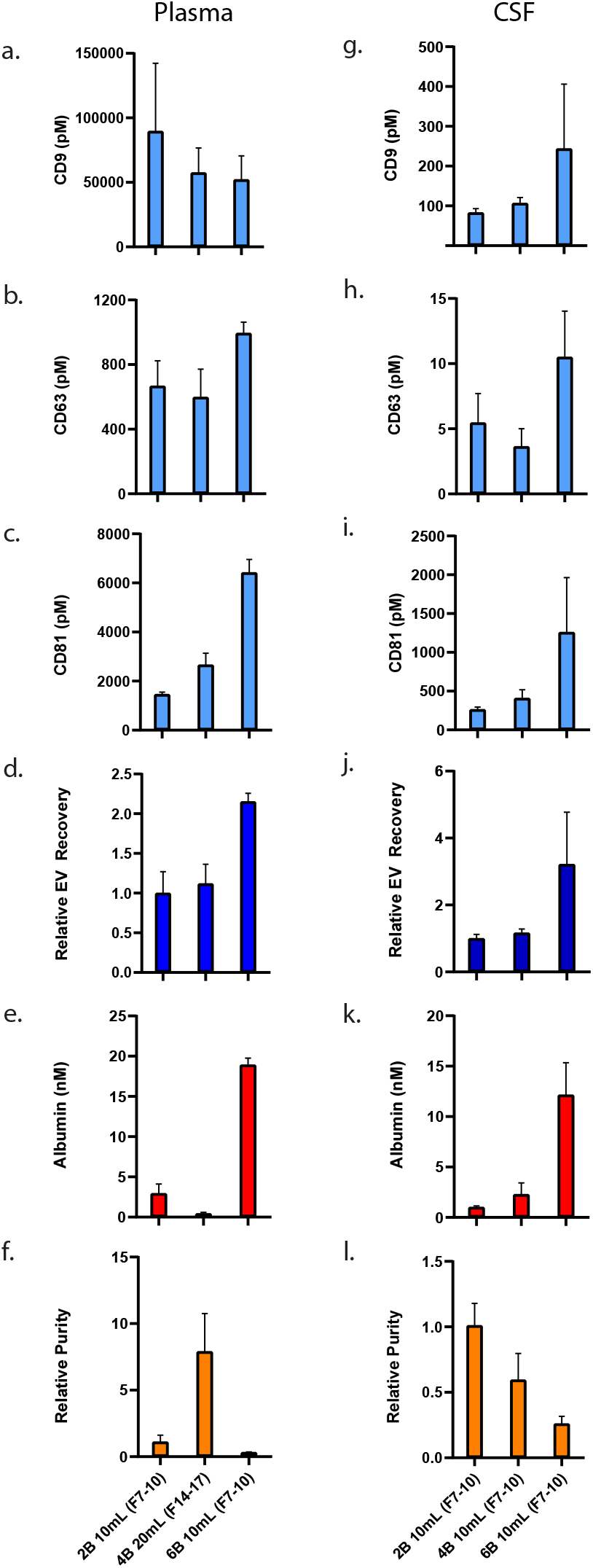
Comparison of top custom SEC methods in plasma and CSF. Error bars represent the standard deviations from four replicates of each column. a-c. Individual tetraspanin yields using different isolation methods from plasma. Relative EV recoveries from plasma were calculated by first normalizing individual tetraspanin values (in pM) in each technique to those of the Sepharose CL-2B 10 mL column (fractions 7-10) and then averaging the three tetraspanin ratios. e. Albumin levels using different EV isolation methods from plasma. EV purity for each method in plasma is calculated as the ratio of the sum of tetraspanin concentrations divided by albumin concentration. g-i. Individual tetraspanin yield using different isolation methods from CSF. Relative EV recoveries in CSF were calculated by first normalizing individual tetraspanin values (in pM) in each technique to those of Sepharose CL-2B 10 mL (fractions 7-10) and then averaging the three tetraspanin ratios. k. Albumin levels using different EV isolation methods from CSF. l. EV purity for each method in CSF is calculated as the ratio of the sum of tetraspanin concentrations divided by albumin concentration.

## Discussion

In this study, we describe a framework for rapidly quantifying relative EV yield and purity across isolation methods, overcoming limitations of other commonly-used methods used for EV analysis. Several techniques, such as Nanoparticle Tracking Analysis (NTA) and other methods developed for analysis of synthetic particles, have been applied to EV detection (3). The utility of these techniques is hindered, however, by an inability to differentiate heterogeneous EVs from other particles with overlapping size, such as lipoproteins or aggregated protein particles. Thus, although previous reports comparing EV isolation methods (23–34) have yielded some useful insights, the lack of reliable EV quantification has made these studies difficult to interpret (2, 5, 35).

The measurement of EV transmembrane proteins overcomes the limitations of EV quantification with particle detection methods. Since the transmembrane proteins CD9, CD63, and CD81 are present on EVs but are not present in lipoproteins or free protein aggregates, these tetraspanins can be used for relative quantification of EVs. Although not every EV necessarily contains a tetraspanin protein, by detecting three different tetraspanins per sample with Simoa, we minimize the chance that we are measuring a rare subset of EVs. In the experiments reported here, we observed a strong correlation of the relative levels of the three tetraspanins in different SEC fractions. Since we compared isolation methods from the same starting sample, we were able to provide a direct quantitative comparison of tetraspanin levels between the different isolation methods.

We used Simoa in this study, which is particularly well suited for EV analysis due to the technology’s high dynamic range, throughput, and sensitivity. This sensitivity is achieved by converting ELISA to a digital readout via immuno-capture and counting of individual protein molecules in a microwell array. We used the commercially-available Quanterix HD-X instrument, but our lab has also developed other digital ELISA methods using commonly-available instrumentation (36–38), which could be similarly applied to EVs. One could also follow a similar approach to the one we present here with traditional ELISA or other protein detection methods, but we find that high sensitivity is often necessary for the low levels of EVs in human biofluids. We have previously shown in a direct comparison (using the same antibodies) that Simoa can detect EV markers in cases where traditional ELISA cannot, such as SEC fractions of CSF (19).

We used Simoa to directly compare yield and purity of commonly used EV isolation methods. To obtain the purest EVs possible (and separate EVs from lipoproteins), it has been demonstrated that it is necessary to combine several techniques sequentially, such as density gradient centrifugation (DGC) and SEC (39, 40). However, techniques such as DGC are not scalable to many samples and therefore not amenable to biomarker studies. Thus, we focused on EV isolation methods that amenable to biomarker studies. After finding that commercial SEC columns compare favorably to ultracentrifugation and ExoQuick precipitation, we compared several resins and column volumes to further improve EV isolation by custom SEC columns.

Our investigation of SEC parameters led us to improved methods for EV isolation; in particular, we found that Sepharose CL-6B, which is seldom used for EV isolation, yields considerably higher levels of EVs than either Sepharose CL-2B, the most commonly used resin (22), or Sepharose CL-4B. We attribute this result to Sepharose CL-6B beads having a smaller average pore size, leading to a lower probability that EVs will enter the beads. As there is a tradeoff between EV yield and albumin contamination, we envision different SEC columns will be suited for different applications. Using a 10 mL Sepharose CL-6B column for EV isolation from plasma or CSF is the best choice for downstream applications where maximum EV yield is needed and where some free protein contamination is not detrimental - for example, analyzing rare EV cargo or when further purification of EVs will be performed (such as immuno-isolation). On the other hand, if isolating EVs from plasma where minimal free protein contamination is desired (for example, in EV protein analysis by Western Blot), a larger 20 mL column with Sepharose CL-4B would yield better results. For CSF, which has much less protein than plasma, 10 mL columns are preferable to 20 mL ones.

By developing a Simoa assay to measure albumin (the most abundant protein in plasma and main contaminant when isolating EVs), we were able to assess the purity of EV preparations with respect to unwanted co-purification of free proteins. Our methods could be expanded to assess other contaminants that are less abundant than albumin but may, nonetheless, be problematic for some applications, such as lipoproteins. Adding a Simoa assay for ApoB100 (or other protein components of lipoproteins) would allow for the assessment of both lipoprotein and free protein contamination in EV isolation methods. Although lipoproteins are difficult to separate from EVs due to their overlapping size profile (12), a recent study demonstrated that a chromatography column combining a cation-exchange resin layer with an SEC resin layer allows for efficient lipoprotein depletion using “dual mode chromatography” (41). Simoa could be used to evaluate and help improve such techniques in the future.

The general experimental framework presented here could be easily applied to evaluate new EV isolation methods in plasma, CSF, or other biological fluids, such as urine or saliva. While we limited our study to human biofluids, similar methods could also be applied to compare EV isolation methods from cell culture media. As sensitivity of EV detection and specificity in differentiating EVs from contaminants are obstacles in all EV studies, we envision that ultrasensitive protein detection with Simoa will be broadly applicable to assessing EV isolation methods for both the study of EV biology and development of EV diagnostics.

## Materials and Methods

### Human Sample Handling

Pre-aliquoted pooled human plasma and CSF samples were ordered from BioIVT. The same pools were used for all main figures throughout the paper in order to ensure comparable analysis of methods. For all EV isolation technique comparisons, one 0.5 mL sample was used for each isolation method. Plasma or CSF was thawed at room temperature. After sample thawing, 100X Protease/Phosphatase Inhibitor Cocktail (Cell Signaling Technology) was added to 1X. The sample was then centrifuged at 2000 × *g* for 10 minutes. The supernatant was subsequently centrifuged through a 0.45 μm Corning Costar SPIN-X centrifuge tube filter (Sigma-Aldrich) at 2000 × *g* for 10 minutes to get rid of any remaining cells or cell debris.

### Simoa Assays

Simoa assays were developed and performed as previously described (19). Capture antibodies were coupled to Carboxylated Paramagnetic Beads from the Simoa Homebrew Assay Development Kit (Quanterix) using EDC chemistry (Thermo Fisher Scientific). Detection antibodies were conjugated to biotin using EZ-Link NHS-PEG4 Biotin (Thermo Fisher Scientific). The following antibodies were used as capture antibodies for tetraspanins: ab195422 (Abcam), MAB5048 (R&D Systems), and ab79559 (Abcam). The following antibodies were used as detector antibodies for tetraspanins: ab58989 (R&D Systems), 556019 (BD) and 349502 (BioLegend). For albumin, DY1455 (R&D Systems) was used as both capture and detector antibody. The following recombinant proteins were used for CD9, CD63, CD81, and albumin: ab152262 (Abcam), TP301733 (Origene), TP317508 (Origene), ab201876 (Abcam).

On-board dilution was performed with 4X dilution for each of the tetraspanins, while manual 20X dilution was used for albumin. All samples were raised to 160 μL per replicate in sample diluent. For tetraspanin assays, samples were incubated with immunocapture beads (25 μL) and biotinylated detection antibody (20 μL) for 35 minutes. Next, six washes were performed, and the beads were resuspended in 100 μL of Streptavidin labeled β-galactosidase (Quanterix) and incubated for 5 minutes. An additional six washes were performed, and the beads were resuspended in 25 μL Resorufin β-D-galactopyranoside (Quanterix) before being loaded into the microwell array on the Quanterix HD-X instrument.

For the albumin assay, samples were incubated first with immunocapture beads (25 μL) for 15 minutes and then washed six times. Subsequently, 100 μL detection antibody was incubated with the beads for 5 minutes. Next, six washes were performed, and the beads were resuspended in 100 μL of Streptavidin labeled β-galactosidase (Quanterix) for a final 5-minute incubation. After an additional six washes, the beads were resuspended in 25 μL Resorufin β-D-galactopyranoside (Quanterix) and then loaded into the microwell array on the Quanterix HD-X instrument.

### Construction of SEC Stand

The custom SEC rack was constructed from a total of 22 pieces using CNC milling tools. The rack is made up of an aluminum frame (silver, Multipurpose 6061 Aluminum, McMaster-Carr) consisting of eight pieces, four sliding plates made from acetal (black, Wear-Resistant Easy-to-Machine Delrin® Acetal Resin, McMaster-Carr), and ten sliding plate grips made from UHMW Polyethylene (white, Slippery UHMW Polyethylene, McMaster-Carr). The rack frame is held together using 20 ¾” screws (McMaster-Carr, 92210A113), 20 ½” screws (McMaster-Carr, 92210A110), 10 0.375” Dowel pins (McMaster-Carr, 90145A470), and 10 0.5625” Dowel pins (McMaster-Carr, 90145A483), and includes 20 spring plungers (McMaster-Carr, 84895A710) that allow the sliding plates to “click” once aligned with the chromatography columns. Details for constructing the rack and SolidWorks files are included in the Supplemental Materials.

### Preparation of Custom SEC Columns

The resins Sepharose CL-2B, Sepharose CL-4B, and Sepharose CL-6B (all from GE Healthcare/Cytiva) were washed in PBS. The volume of resin was washed with an equal volume of PBS in a glass container and then placed at 4 °C in order to let the resin settle completely (several hours or overnight). The PBS was then poured off, and an equal volume of PBS was again added two more times for a total of three washes. Columns were prepared fresh on the day of use. Washed resin was poured into an Econo-Pac Chromatography column (Bio-Rad) to bring the bed volume (the resin without liquid) to 10 mL or 20 mL. When the desired amount of resin filled the column and the liquid dripped through, the top frit was immediately placed at the top of the resin without compressing the resin. PBS was then added again before sample addition.

### Collection of Size Exclusion Chromatography Fractions

Once prepared, all columns were washed with at least 20 mL of PBS in the column. Immediately before sample addition, the column was allowed to fully drip out and, after last drop of PBS, sample (filtered plasma or CSF) was added to the column. As soon as sample was added, 0.5 mL fractions were collected in individual tubes. As soon as the plasma or CSF completely entered the column (below the frit), PBS was added to the top of column 1 mL at a time. Fraction numbers correspond to 0.5mL increments collected as soon as sample is added. For Izon and 10 mL columns, fractions 6-21 were collected (since first few fractions correspond to void volume). For 20 mL columns, fractions 12-27 were collected (since void volume is larger for 20 mL columns than 10 mL columns). For Figure 6, only fractions 7-10 were collected.

### Ultracentrifugation

Samples of filtered 0.5 mL plasma or CSF were added to 3.5 mL Open-Top Thickwall Polycarbonate ultracentrifuge tubes (Beckman Coulter), and PBS was added to fill tubes to the top. Samples were ultracentrifuged at 120,000 × *g* for 90 minutes at 4 °C in an Optima XPN-80 ultracentrifuge (Beckman Coulter) using a SW55 Ti swinging-bucket rotor (Beckman Coulter). Afterwards, all supernatant was aspirated. Pellets were resuspended in PBS for the “Ultracentrifugation” condition. For the “Ultracentrifugation with wash” condition, the ultracentrifuge tubes were filled to the top with PBS, and samples were ultracentrifuged again at 120,000 × *g* for 90 minutes. Supernatant was then aspirated, and pellets were resuspended in 500 uL PBS. For all ultracentrifugation samples, isolation was performed on two separate days and then resulting Simoa values were averaged.

### ExoQuick & ExoQuick ULTRA

Samples of plasma or CSF were mixed with ExoQuick Exosome Precipitation Solution (System Biosciences) or ExoQuick ULTRA EV Isolation Kit for Serum and Plasma (System Biosciences), and protocols were performed according to manufacturer’s instructions. For ExoQuick, 0.5 mL of plasma or CSF was mixed with 126 uL of ExoQuick and incubated at 4 °C for 30 minutes, followed by centrifugation at 1500 × *g* for 30 minutes. Supernatant was removed, and samples were centrifuged at 1500 × *g* for an additional 5 minutes. Residual supernatant was removed, and pellets were resuspended in 500 uL PBS. For ExoQuick ULTRA, 250 uL of plasma or CSF was used in accordance with instructions, and Simoa values were corrected by multiplying by two to match 0.5 mL volume used for other samples. For each sample, 500 uL of EVs was eluted per column. For all precipitations, isolation was performed on two separate days and then resulting Simoa values were averaged.

## Supporting information

SI SEC Stand Assembly Instructions

## Author Contributions

DT and MN conceived of study and designed experiments. DT, MN, WT, JHL, and RL performed experiments. DT, MN, and DRW analyzed data and wrote the manuscript with input from all authors. GMC and DRW supervised the study and provided funding support.

## Acknowledgements

The authors acknowledge David Kalish for help designing and making the SEC stand and Emma Kowal for help with illustrations and comments on the manuscript. Funding for this study was provided by the Chan Zuckerberg Initiative (CZI) Neurodegeneration Challenge Network (NDCN) and Good Ventures Foundation. Schematics were created with BioRender.com

---

**Supplementary Figure 1:**
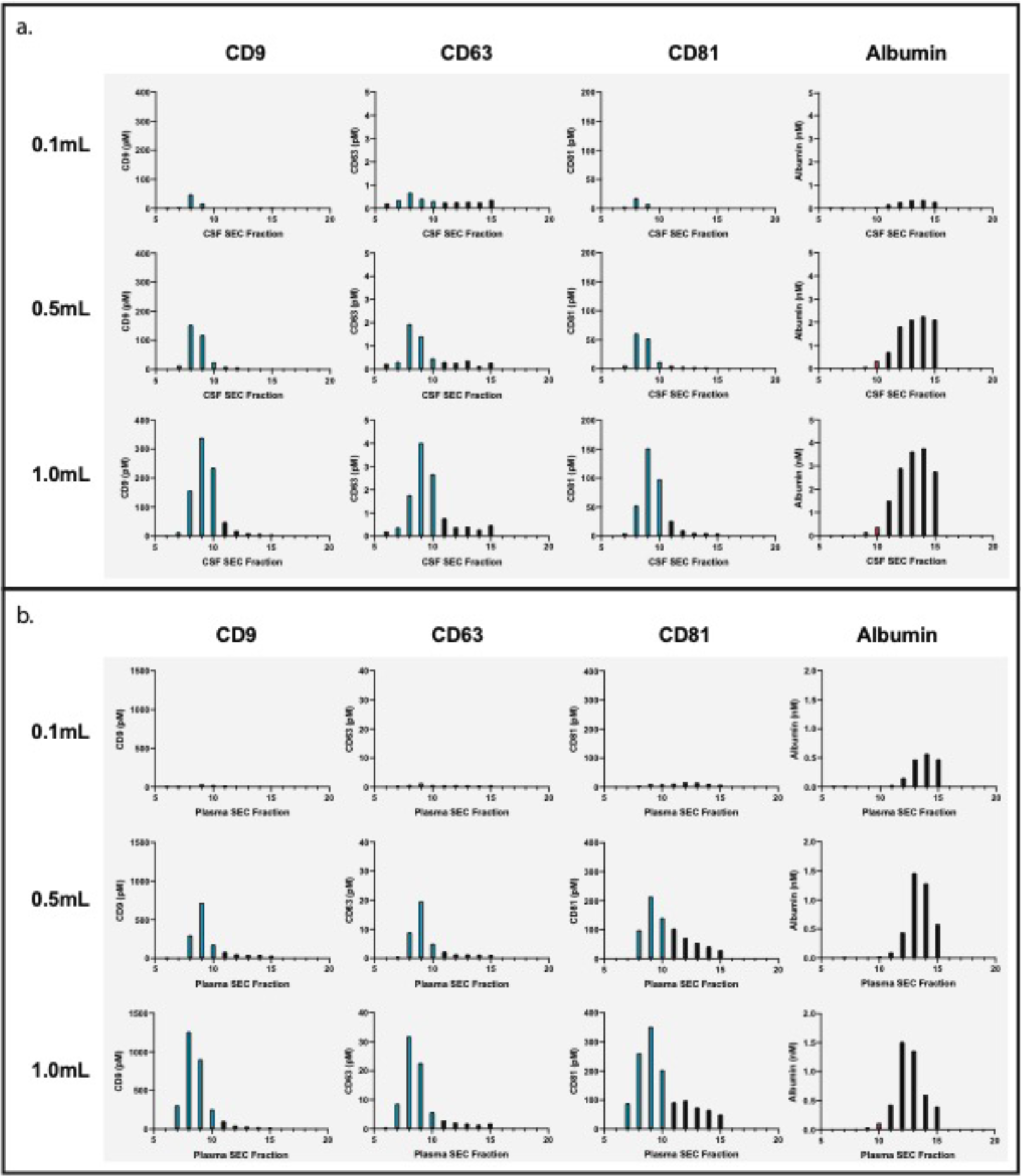
Effect of CSF and plasma sample volume on SEC. a. Effect of sample volume on EV recovery and purity by SEC for 0.1 mL (top), 0.5 mL (middle) and 1.0 mL (bottom) samples of CSF. Simoa was performed to determine levels of CD9, CD63, CD81 and albumin after fractionating different volumes of CSF by SEC using a 10 mL Sepharose CL-6B column. b. Effect of sample volume on EV recovery and purity by SEC for 0.1 mL (top), 0.5 mL (middle) and 1.0 mL (bottom) samples of plasma. Simoa was performed to determine levels of CD9, CD63, CD81 and albumin after fractionating different volumes of plasma by SEC using a 10 mL Sepharose CL-6B column.

**Supplementary Figure 2:**
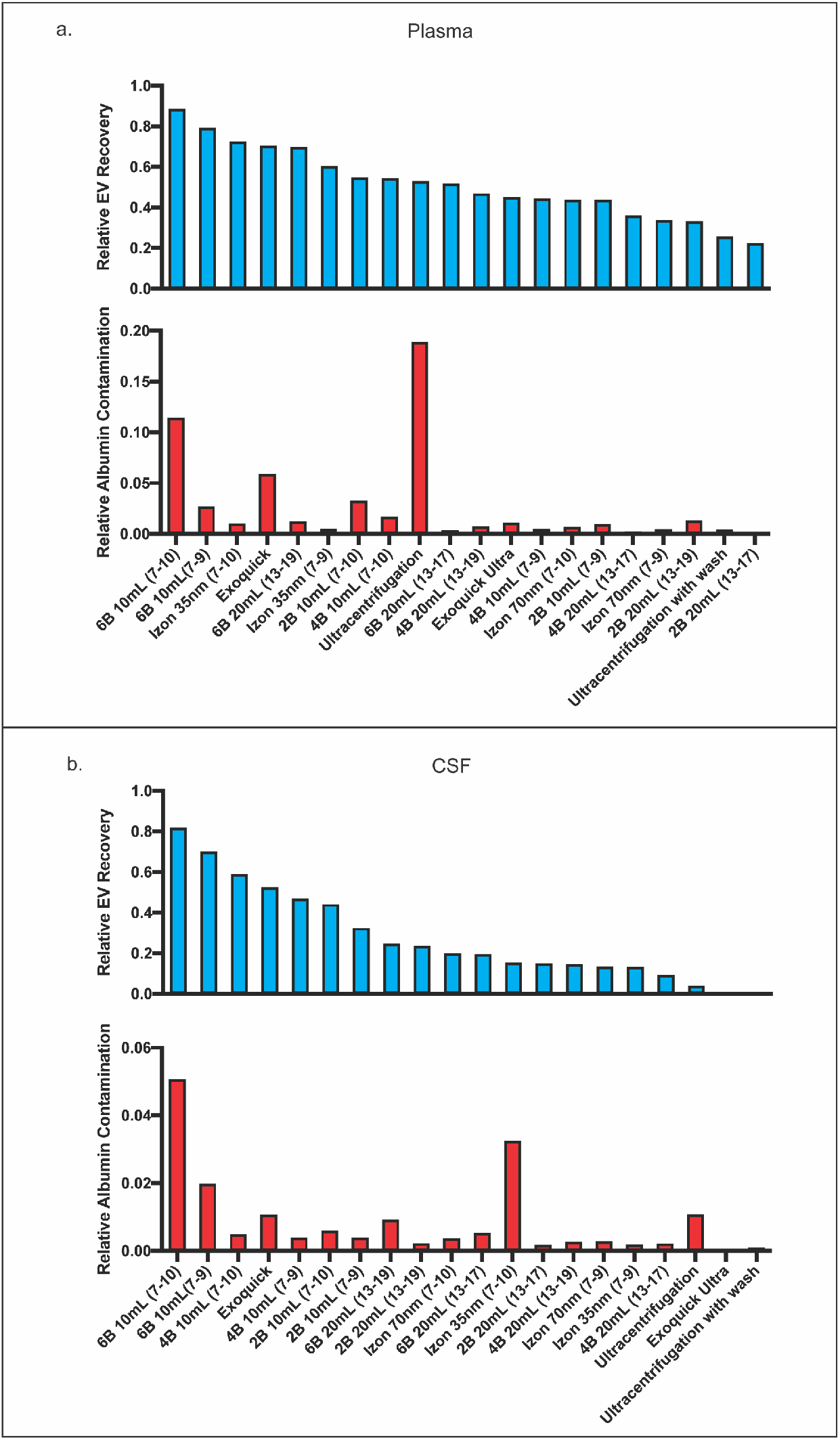
Comparison of EV recovery and albumin contamination across all tested methods in plasma and CSF. a. Comparison of plasma EV recovery (top) and albumin contamination (bottom) across all tested methods ranked by EV recovery. Relative EV recoveries were calculated by individually normalizing each tetraspanin to the sum of the tetraspanins in all fractions (6–21) in the 10mL Sepharose CL-6B condition. The three tetraspanin percentages were then averaged to calculate the relative EV recoveries. Similarly, albumin for each condition was calculated as a fraction of the albumin found in all fractions (6–21) in the 10mL Sepharose CL-6B condition. b. Comparison of CSF EV recovery (top) and albumin contamination (bottom) across all tested methods ranked by EV recovery. Relative EV recovery was calculated by individually normalizing each tetraspanin to the sum of that tetraspanin in all fractions (6–21) in the 10mL Sepharose CL-6B condition. The three tetraspanin percentages were then averaged to calculate the relative EV recovery. Similarly, albumin for each condition was calculated as a fraction of the albumin found in all fractions (6–21) in the 10mL Sepharose CL-6B condition.

## References

1. Hirshman BR, Kras RT, Akers JC, Carter BS, Chen CC. Extracellular Vesicles in Molecular Diagnostics: An Overview with a Focus on CNS Diseases. Advances in clinical chemistry. 2016;76:37–53.

2. Tkach M, Kowal J, Thery C. Why the need and how to approach the functional diversity of extracellular vesicles. Philosophical transactions of the Royal Society of London Series B, Biological sciences. 2018;373(1737).

3. Hartjes TA, Mytnyk S, Jenster GW, van Steijn V, van Royen ME. Extracellular Vesicle Quantification and Characterization: Common Methods and Emerging Approaches. Bioengineering (Basel, Switzerland). 2019;6(1).

4. Shao H, Im H, Castro CM, Breakefield X, Weissleder R, Lee H. New Technologies for Analysis of Extracellular Vesicles. Chemical reviews. 2018;118(4):1917–50.

5. Coumans FAW, Brisson AR, Buzas EI, Dignat-George F, Drees EEE, El-Andaloussi S, et al. Methodological Guidelines to Study Extracellular Vesicles. Circulation research. 2017;120(10):1632–48.

6. Konoshenko MY, Lekchnov EA, Vlassov AV, Laktionov PP, Isolation of Extracellular Vesicles: General Methodologies and Latest Trends. BioMed research international. 2018;2018:8545347.

7. Théry C, Witwer KW, Aikawa E, Alcaraz MJ, Anderson JD, Andriantsitohaina R, et al. Minimal information for studies of extracellular vesicles 2018 (MISEV2018): a position statement of the International Society for Extracellular Vesicles and update of the MISEV2014 guidelines. Journal of extracellular vesicles. 2018;7(1):1535750.

8. Sodar BW, Kittel A, Paloczi K, Vukman KV, Osteikoetxea X, Szabo-Taylor K, et al. Low-density lipoprotein mimics blood plasma-derived exosomes and microvesicles during isolation and detection. Scientific reports. 2016;6:24316.

9. Welton JL, Webber JP, Botos LA, Jones M, Clayton A, Ready-made chromatography columns for extracellular vesicle isolation from plasma. Journal of extracellular vesicles. 2015;4:27269.

10. Lee YXF, Johansson H, Wood MJA, El Andaloussi S, Considerations and Implications in the Purification of Extracellular Vesicles - A Cautionary Tale. Front Neurosci. 2019;13:1067.

11. Webber J, Clayton A. How pure are your vesicles? Journal of extracellular vesicles. 2013;2.

12. Simonsen JB. What Are We Looking At? Extracellular Vesicles. Lipoproteins, or Both? Circulation research. 2017;121(8):920–2.

13. Osteikoetxea X, Balogh A, Szabo-Taylor K, Nemeth A, Szabo TG, Paloczi K, et al. Improved characterization of EV preparations based on protein to lipid ratio and lipid properties. PloS one. 2015;10(3):e0121184.

14. Visnovitz T, Osteikoetxea X, Sodar BW, Mihaly J, Lorincz P, Vukman KV, et al. An improved 96 well plate format lipid quantification assay for standardisation of experiments with extracellular vesicles. Journal of extracellular vesicles. 2019;8(1):1565263.

15. Lucchetti D, Battaglia A, Ricciardi-Tenore C, Colella F, Perelli L, De Maria R, et al. Measuring Extracellular Vesicles by Conventional Flow Cytometry: Dream or Reality? International journal of molecular sciences. 2020;21(17).

16. Kuiper M, van de Nes A, Nieuwland R, Varga Z, van der Pol E, Reliable measurements of extracellular vesicles by clinical flow cytometry. Am J Reprod Immunol. 2021;85(2):e13350.

17. Welsh JA, Van Der Pol E, Arkesteijn GJA, Bremer M, Brisson A, Coumans F, et al. MIFlowCyt-EV: a framework for standardized reporting of extracellular vesicle flow cytometry experiments. Journal of extracellular vesicles. 2020;9(1):1713526.

18. Coumans FAW, Gool EL, Nieuwland R. Bulk immunoassays for analysis of extracellular vesicles. Platelets. 2017;28(3):242–8.

19. Norman M, Ter-Ovanesyan D, Trieu W, Lazarovits R, Kowal EJK, Lee JH, et al. L1CAM is not Associated with Extracellular Vesicles in Human Cerebrospinal Fluid or Plasma. bioRxiv. 2020:2020.08.12.247833.

20. Cohen L, Walt DR. Highly Sensitive and Multiplexed Protein Measurements. Chemical reviews. 2019;119(1):293–321.

21. Boing AN, van der Pol E, Grootemaat AE, Coumans FA, Sturk A, Nieuwland R. Single-step isolation of extracellular vesicles by size-exclusion chromatography. Journal of extracellular vesicles. 2014;3.

22. Monguio-Tortajada M, Galvez-Monton C, Bayes-Genis A, Roura S, Borras FE. Extracellular vesicle isolation methods: rising impact of size-exclusion chromatography. Cellular and molecular life sciences : CMLS. 2019;76(12):2369–82.

23. Lobb RJ, Becker M, Wen SW, Wong CS, Wiegmans AP, Leimgruber A, et al. Optimized exosome isolation protocol for cell culture supernatant and human plasma. Journal of extracellular vesicles. 2015;4:27031.

24. Helwa I, Cai J, Drewry MD, Zimmerman A, Dinkins MB, Khaled ML, et al. A Comparative Study of Serum Exosome Isolation Using Differential Ultracentrifugation and Three Commercial Reagents. PloS one. 2017;12(1):e0170628.

25. Baranyai T, Herczeg K, Onodi Z, Voszka I, Modos K, Marton N, et al. Isolation of Exosomes from Blood Plasma: Qualitative and Quantitative Comparison of Ultracentrifugation and Size Exclusion Chromatography Methods. PloS one. 2015;10(12):e0145686.

26. Soares Martins T, Catita J, Martins Rosa I, O ABdCES, Henriques AG. Exosome isolation from distinct biofluids using precipitation and column-based approaches. PloS one. 2018;13(6):e0198820.

27. An M, Wu J, Zhu J, Lubman DM. Comparison of an Optimized Ultracentrifugation Method versus Size-Exclusion Chromatography for Isolation of Exosomes from Human Serum. Journal of proteome research. 2018;17(10):3599–605.

28. Stranska R, Gysbrechts L, Wouters J, Vermeersch P, Bloch K, Dierickx D, et al. Comparison of membrane affinity-based method with size-exclusion chromatography for isolation of exosome-like vesicles from human plasma. Journal of translational medicine. 2018;16(1):1.

29. Diaz G, Bridges C, Lucas M, Cheng Y, Schorey JS, Dobos KM, et al. Protein Digestion, Ultrafiltration, and Size Exclusion Chromatography to Optimize the Isolation of Exosomes from Human Blood Plasma and Serum. Journal of visualized experiments : JoVE. 2018(134).

30. Kalra H, Adda CG, Liem M, Ang CS, Mechler A, Simpson RJ, et al. Comparative proteomics evaluation of plasma exosome isolation techniques and assessment of the stability of exosomes in normal human blood plasma. Proteomics. 2013;13(22):3354–64.

31. Serrano-Pertierra E, Oliveira-Rodriguez M, Rivas M, Oliva P, Villafani J, Navarro A, et al. Characterization of Plasma-Derived Extracellular Vesicles Isolated by Different Methods: A Comparison Study. Bioengineering (Basel, Switzerland). 2019;6(1).

32. Gamez-Valero A, Monguio-Tortajada M, Carreras-Planella L, Franquesa M, Beyer K, Borras FE. Size-Exclusion Chromatography-based isolation minimally alters Extracellular Vesicles’ characteristics compared to precipitating agents. Scientific reports. 2016;6:33641.

33. Takov K, Yellon DM, Davidson SM. Comparison of small extracellular vesicles isolated from plasma by ultracentrifugation or size-exclusion chromatography: yield, purity and functional potential. Journal of extracellular vesicles. 2019;8(1):1560809.

34. Brennan K, Martin K, FitzGerald SP, O’Sullivan J, Wu Y, Blanco A, et al. A comparison of methods for the isolation and separation of extracellular vesicles from protein and lipid particles in human serum. Scientific reports. 2020;10(1):1039.

35. Ludwig N, Whiteside TL. Reichert TE, Challenges in Exosome Isolation and Analysis in Health and Disease. International journal of molecular sciences. 2019;20(19).

36. Cohen L, Cui N, Cai Y, Garden PM, Li X, Weitz DA, et al. Single Molecule Protein Detection with Attomolar Sensitivity Using Droplet Digital Enzyme-Linked Immunosorbent Assay. ACS nano. 2020;14(8):9491–501.

37. Maley AM, Garden PM, Walt DR. Simplified Digital Enzyme-Linked Immunosorbent Assay Using Tyramide Signal Amplification and Fibrin Hydrogels. ACS Sens. 2020.

38. Wu C, Garden PM, Walt DR. Ultrasensitive Detection of Attomolar Protein Concentrations by Dropcast Single Molecule Assays. J Am Chem Soc. 2020;142(28):12314–23.

39. Karimi N, Cvjetkovic A, Jang SC, Crescitelli R, Hosseinpour Feizi MA, Nieuwland R, et al. Detailed analysis of the plasma extracellular vesicle proteome after separation from lipoproteins. Cellular and molecular life sciences : CMLS. 2018;75(15):2873–86.

40. Zhang X, Borg EGF, Liaci AM, Vos HR, Stoorvogel W. A novel three step protocol to isolate extracellular vesicles from plasma or cell culture medium with both high yield and purity. Journal of extracellular vesicles. 2020;9(1):1791450.

41. Van Deun J, Jo A, Li H, Lin HY, Weissleder R, Im H, et al. Integrated Dual-Mode Chromatography to Enrich Extracellular Vesicles from Plasma. Advanced biosystems. 2020;4(12):e1900310.

