## Supplementary material for "Framework for Rapid Comparison of Extracellular Vesicle Isolation Methods": SI SEC Stand Assembly Instructions: SI SEC Stand Assembly Instructions.pdf

Supplemental File 1: Instructions for Constructing Custom SEC Stand

To construct the rack, become familiar with the general rack structure by reviewing the SolidWorks assemblies included in the Supplemental Information CAD files . A summary of parts required to assemble the rack and pictures of each part are included in the tables below. Begin the rack assembly by constructing five Slots (use the Slot Assembly file as reference while constructing this piece). When putting together the Slots, begin by attaching four Spring Screws to the Slot Bottom piece. After attaching the Spring Screws, combine the Slot Bottom, Slot Middle, and Slot Top pieces using 9/16” Dowel Pins (90145A483). This assembly should then be screwed into a Slot Fin using ¾” screws (92210A113), completing one Slot piece. Repeat this process for each of the five Slots.

After assembling the Slots, begin assembling the Rack Frame (use the Rack Frame assembly file as reference). Start by attaching one Rack Side piece to the Rack Bottom piece using ½” screws (for the remainder of this section, all screws should be ½” screws, 92210A110). Next, attach one side of the Falcon Tube Tip Plate, Back Bar, Column Tip Plate, and Column Barrel Plate to the same Rack Side piece. Do not attach the Falcon Tube Barrel Plate until after the next step. Now, attach the five Slots from above to the Back Bar using two 3/8” Dowel Pins (90145A470) and a screw. Finally, attach the Falcon Tube Barrel Plate to the Rack Side piece, and complete the rack by screwing in the remaining Rack Side piece. The chromatography column rack is now ready for use, and the 2mL Microtube Plates can be inserted in between the slots.

| Piece | Quantity | Catalog Number | Comments |
|---|---|---|---|
| Rack Side | 2 | CNC Milled, Aluminum | |
| Rack Bottom | 1 | CNC Milled, Aluminum | |
| Falcon Tube Tip Plate | 1 | CNC Milled, Aluminum | |
| Back Bar | 1 | CNC Milled, Aluminum | |
| Column Tip Plate | 1 | CNC Milled, Aluminum | |
| Column Barrel Plate | 1 | CNC Milled, Aluminum | |
| Falcon Tube Barrel Plate | 1 | CNC Milled, Aluminum | |
| 2 mL Microtube Plate | 4 | CNC Milled, Delrin | |
| Slot Bottom | 5 | CNC Milled, UHMW Polyethylene | |
| Slot Middle | 5 | CNC Milled, UHMW Polyethylene | |
| Slot Top | 5 | CNC Milled, Aluminum | |
| Slot Fin | 5 | CNC Milled, Aluminum | |
| Spring Screw | 20 | 84895A710 | .16in x .16in ss18-8 ball-nose spring plunger |
| 3/8” Dowel Pin | 10 | 90145A470 | .125in x .375in ss18-8 dowel pin |
| 9/16” Dowel Pin | 5 | 90145A483 | .125in x .5625in ss18-8 dowel pin |
| ½” Screw | 20 | 92210A110 | 4-40 x .5in ss18-8 c-sink screw |
| ¾” Screw | 20 | 92210A113 | 4-40 x .75in ss18-8 c-sink screw |

| Piece | Picture |
|---|---|
| Rack Assembly | 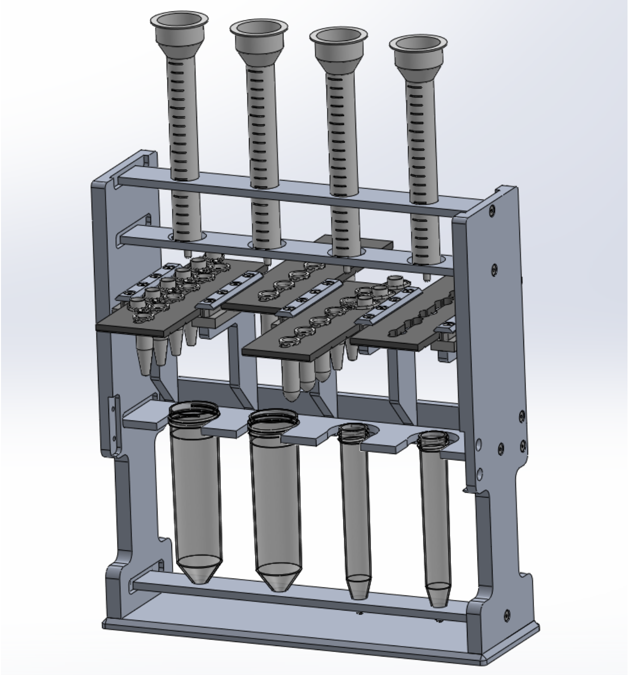 |
| Slot Assembly | 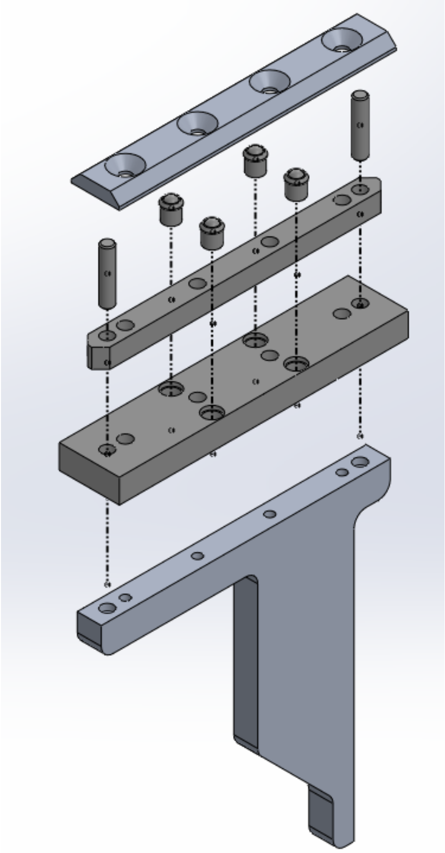 |

| | |
|---|---|
| Rack Side | 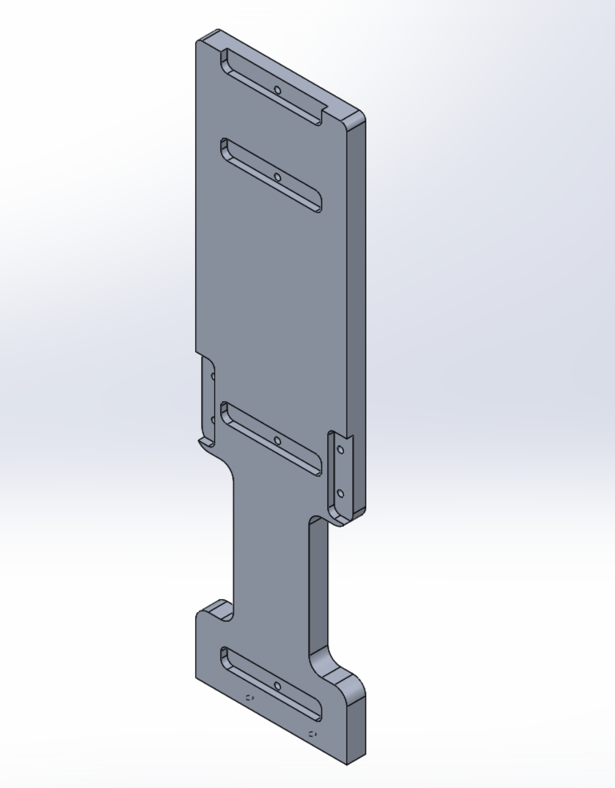 |
| Rack Bottom | 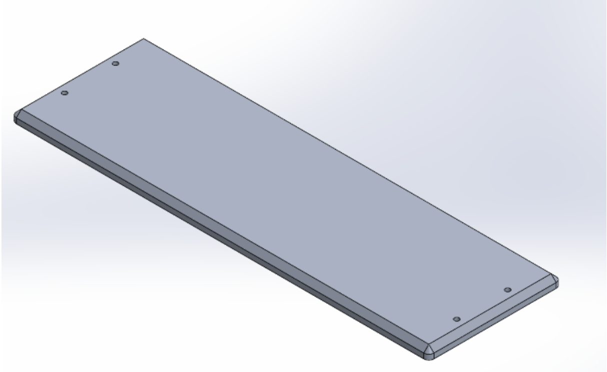 |
| Falcon Tube Tip Plate | 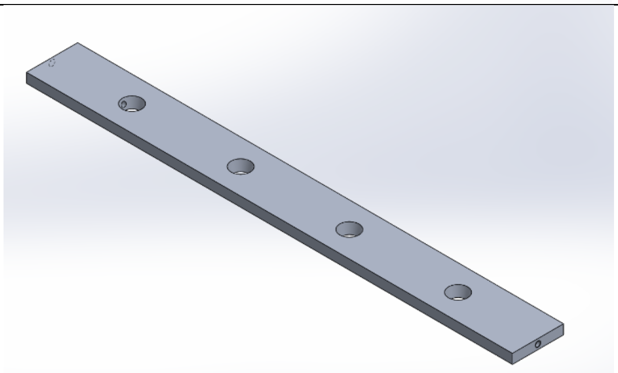 |

| | |
|---|---|
| Back Bar | 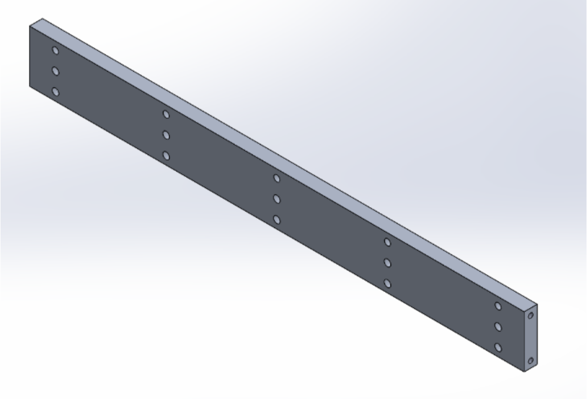 |
| Column Tip Plate | 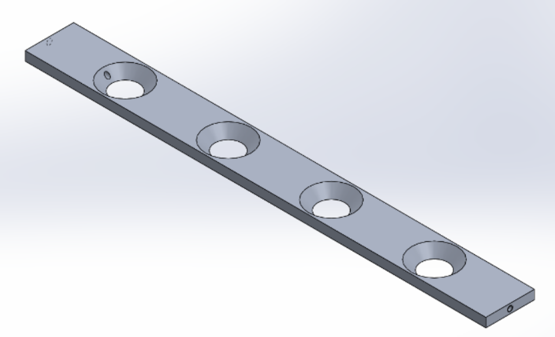 |
| Column Barrel Plate | 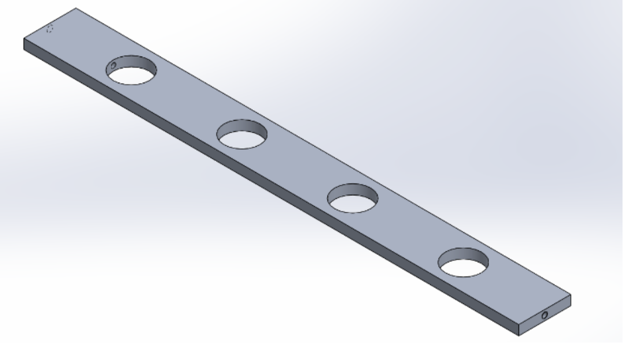 |
| Falcon Tube Barrel Plate | 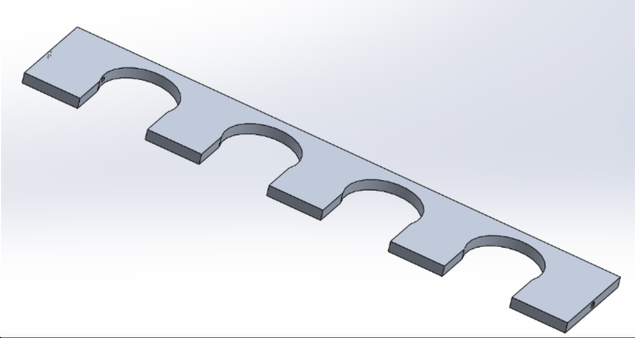 |

| | |
|---|---|
| 2 mL Microtube Plate | 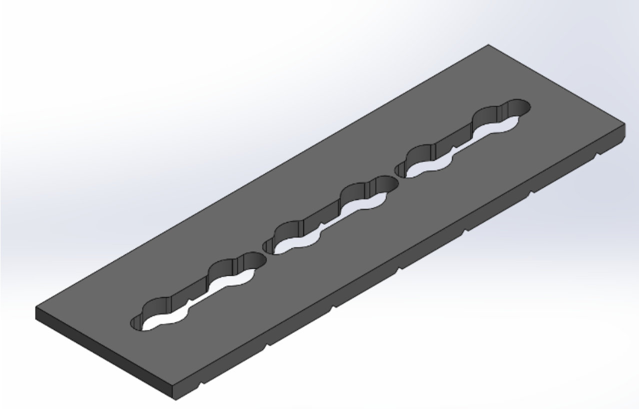 |
| Slot Bottom | 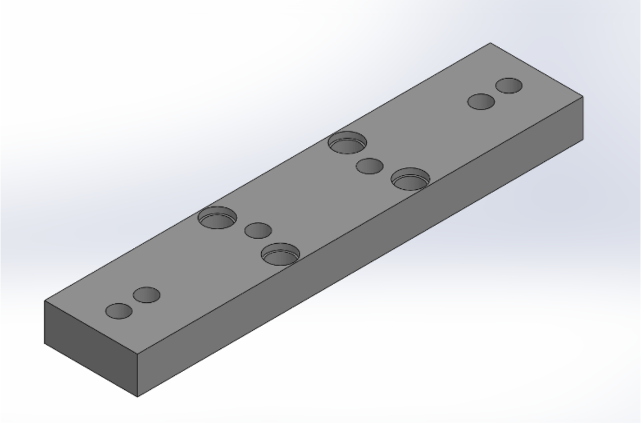 |
| Slot Middle | 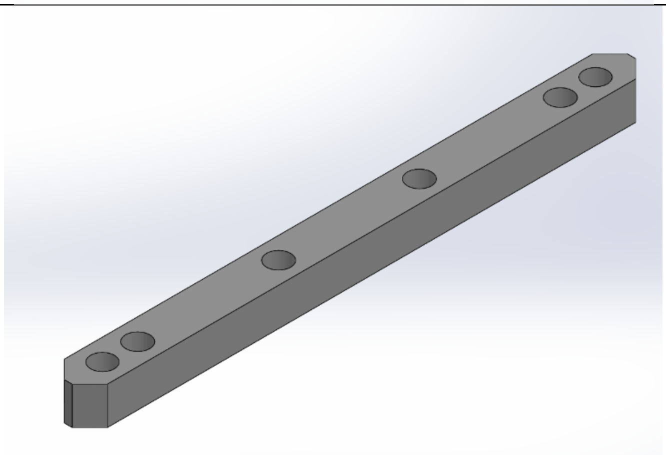 |
| Slot Top | 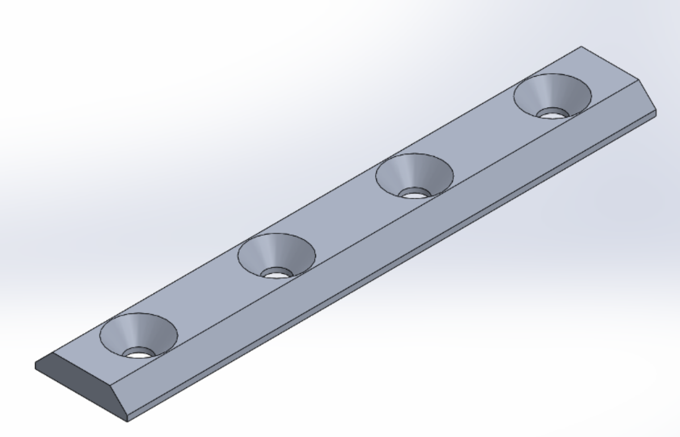 |

| | |
|---|---|
| Slot Fin | 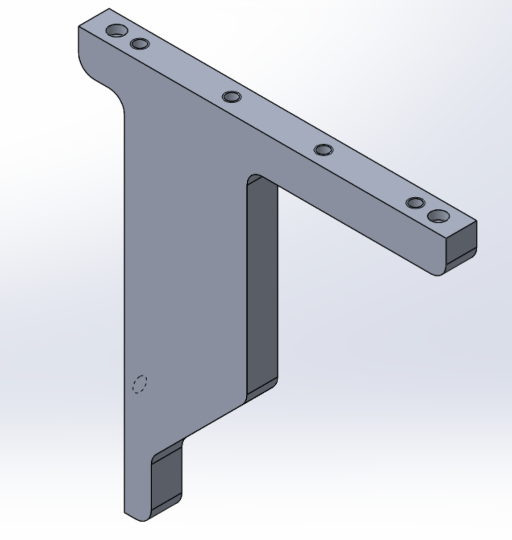 |
| Spring Screw | 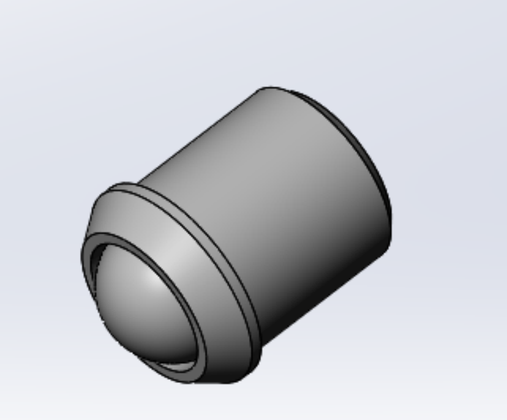 |
| 3/8” Dowel Pin | 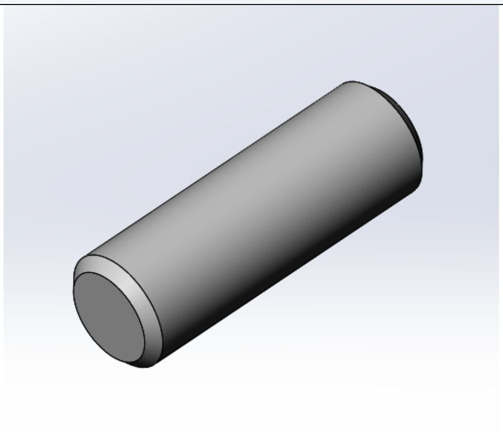 |

| | |
|---|---|
| 9/16" Dowel Pin | 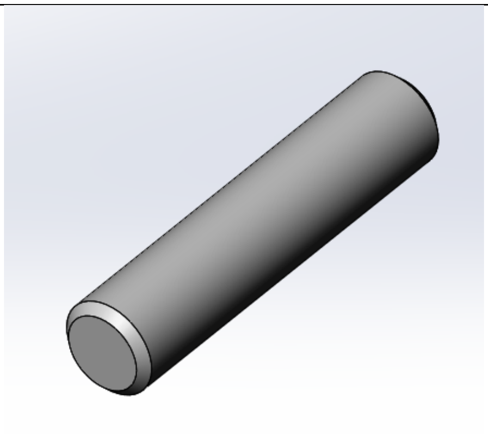 |
| ½" Screw | 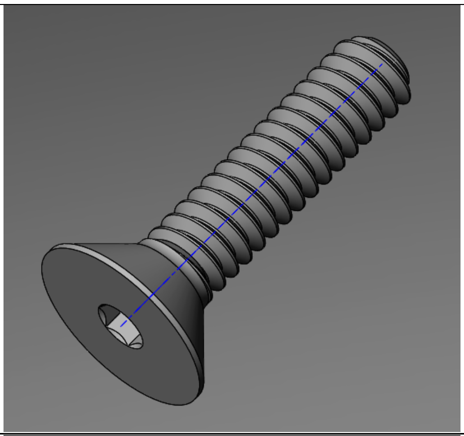 |
| ¾" Screw | 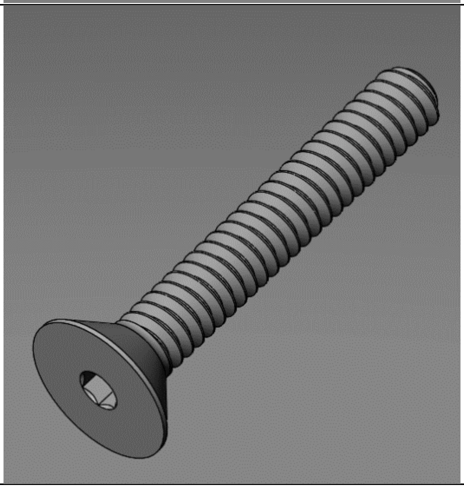 |
